# Citizen science informs demand-driven breeding of opportunity crops

**DOI:** 10.1101/2024.07.31.605211

**Authors:** Rachel C. Voss, Kauê de Sousa, Sognigbé N’Danikou, Abdul Shango, Lys Amavi Aglinglo, Marie-Angélique Laporte, Eric C. Legba, Aristide Carlos Houdegbe, Danfing dit Youssouf Diarra, Aminata Dolo, Amadou Sidibe, Colette Ouedraogo, Harouna Coulibaly, Enoch G. Achigan-Dako, Aishi Kileo, Dickson Malulu, Zamira Matumbo, Fekadu Dinssa, Joost van Heerwaarden, Jacob van Etten, Amritbir Riar, Maarten van Zonneveld

## Abstract

**CONTEXT:** Opportunity crops, also known as neglected and underutilized species (NUS), offer benefits to diversify food systems with nutritious and climate-resilient foods. A major limitation to incorporate these crops in farming systems is the lack of improved varieties impedes farmers accessing quality planting material of these crops.

**OBJECTIVES:** The study explored how citizen science methods can support demand-driven breeding and seed production of opportunity crops using leafy amaranth – a nutritious and hardy vegetable- as a case study. The study identified farmer preferences and market segments, with particular attention to gender and social differentiation.

**METHODS:** We used the tricot approach to conduct participatory on-farm trials of 14 varieties with 2,063 farmers from Benin, Mali, and Tanzania. We then analyzed farmer trait and varietal preferences in aggregate and among segments of farmers, generated using cluster analysis.

**RESULTS:** Farmers’ overall preferences for amaranth varieties was driven principally by plant survival, yield, leaf size, taste, and marketability. Distinct farmer segments (older women generalists, young women specialists, older men generalists, and young men specialists) preferred different varieties depending on gender, business-orientation.

**DISCUSSION AND CONCLUSION:** The farmer segments identified here, along with their unique variety preferences provide valuable information for breeders and seed enterprises, and support demand-driven amaranth breeding and seed system development. We specifically noted the need for breeding programs to understand the preferences of young amaranth specialists, both men and women, and to explore organoleptic and market-related properties of amaranth.

**SIGNIFICANCE:** Our findings on differentiated producer preferences will support scaling seed supply of amaranth in Africa to diversify farming systems with a climate-resilient and nutritious crop. The methods used and lessons learned from our citizen science exercise can be applied to enhance breeding and seed supply of other opportunity crops that are underutilized in Africa or other continents.

**Highlights:**

- Farmer citizen science methods reveal seed market insights for opportunity crops
- Amaranth variety preferences vary widely across gender, age, and countries
- Young amaranth producers focus on market and sensory traits compared to old producers
- Large-scale farmer feedback guides opportunity crop breeding for diverse segments

## Introduction

Supporting diversified farming systems that incorporate fruits and vegetables is an important response to challenges of malnutrition and climate vulnerability in Africa (Covic & Hendriks, 2016; FAO et al., 2021; Harris et al., 2022; Keatinge et al., 2011; von Grebmer et al., 2014). Of particular interest are opportunity crops or neglected and underutilized crop species (NUS), which include native and indigenized vegetables (Mwadzingeni et al., 2021; Van Zonneveld et al., 2023). These species are hailed for their high levels of vitamins, minerals, antioxidants, and dietary fibers (Aworh, 2018; Kamga et al., 2013; Odhav et al., 2007; Yang & Keding, 2009), their contributions to agrobiodiversity and climate resilience (Harris et al., 2022; Mwadzingeni et al., 2021; Slabbert et al., 2004; Van Zonneveld et al., 2023), and their suitability for African smallholder systems under a changing climate (Aworh, 2018; Mwadzingeni et al., 2021; Schreinemachers et al., 2018; van Zonneveld et al., 2023).

Although NUS have long been part of diets in Africa, vegetable consumption in Africa is among the lowest in the world and has been relatively static over time (Afari-Sefa et al., 2012; Afshin et al., 2019; Kalmpourtzidou et al., 2020; Schreinemachers et al., 2021). Vegetable supply is generally insufficient to meet dietary recommendations (Kalmpourtzidou et al., 2020), despite sales of NUS throughout sub-Saharan Africa (Weinberger & Pichop, 2009). Evidence suggests that NUS supply is insufficient to meet year-round demand (Okello et al., 2015; Tatsvarei & Rukasha, 2022), which appears to be increasing among growing urban and peri-urban populations (Dinssa et al., 2016; Karanja et al., 2012; Okello et al., 2015; Tatsvarei & Rukasha, 2022). Supporting expanded production of climate-resilient NUS is therefore a promising means to address nutrition challenges and potentially boost smallholder incomes.

Bolstering NUS production and consumption likely requires a range of interventions, including awareness-raising and demand creation among consumers, reforms to policies and subsidy programs to support NUS cultivation and agronomic advances to improve production, infrastructure development related to post-harvest handling, and expanded breeding to ensure NUS meet producer and consumer needs (McMullin et al., 2021). These intervention points emerge from the many factors underlying the underutilization of NUS, including social stigma around NUS consumption and the deprioritization of NUS in policies, research, and development relative to staple crops (Kansiime et al., 2018; Keatinge et al., 2011, 2015; McMullin et al., 2021; Schreinemachers et al., 2018).

We focus here on breeding of neglected vegetables and access to quality seed as critical components of expanded production and consumption. Farmers need access to seed that meets their needs, priorities, and constraints, aligns with consumer demand, and supports their adaptation to climate stresses (Kansiime & Mastenbroek, 2016). However, limited access to quality planting material often undermines the success of NUS interventions (McMullin et al., 2021). While informal seed systems are often the most accessible and affordable for smallholder farmers (Afari-Sefa et al., 2012; Keatinge et al., 2015; McGuire & Sperling, 2016), informed varietal selection is not always possible, and the quality of seed is a frequent concern—including for NUS (Ayenan et al., 2021). Farmers’ current access to improved NUS varieties of vegetables is largely through seed kits distributed by the World Vegetable Center (WorldVeg) (N’Danikou et al., 2022). However, limited commercial offerings of improved NUS varieties leave many farmers to cultivate NUS varieties with lower yield potential or those at relative risk from climate change, pests, and diseases (Adebooye et al., 2005; Mwadzingeni et al., 2021; Schreinemachers et al., 2018).

Breeding of appropriate varieties underpins improved seed access, whether through formal or informal seed systems. Increased attention to breeding of neglected vegetables has the potential to generate numerous biophysical and nutritional benefits, including improved yields, pest and disease resistance, drought and heat tolerance, and high micronutrient content (Mwadzingeni et al., 2021). However, breeding of neglected vegetables has been overlooked, as the historic focus has been on staple crops in the interest of combating caloric deficiencies (Mwadzingeni et al., 2021; Nabuuma et al., 2022; Santpoort, 2020). At present, farmer preferences and market segments for most NUS are poorly understood, and African seed companies’ capacity for vegetable breeding and seed production is limited (Afari-Sefa et al., 2012). As such, breeders and seed enterprises that might be interested in expanding NUS varietal offerings have little market intelligence to guide them.

Furthermore, varietal release processes and evaluation criteria (including Value for Cultivation and Use-VCU), through which newly developed lines are tested and released for commercial production, were designed principally for cereal crops. As such, they do not always measure characteristics of vegetables that are important to producers and consumers, such as color and shape, long seasonality, shelf life, texture and taste (Afari-Sefa et al., 2012; Keatinge et al., 2015; Schreinemachers et al., 2021; Turner & Bishaw, 2016). As a result, relevant traits for agroecological suitability, stress tolerance, and yield have not been well identified or characterized for many NUS (Dinssa et al., 2016). This means that data to inform critical trade-offs in breeding between yield, abiotic and biotic stress tolerance, nutrition, commercially-relevant characteristics, and other traits are not widely available (Afari-Sefa et al., 2012).

Several gene banks at national and international levels, including those hosted by WorldVeg, hold substantial collections of NUS genetic materials. One challenge has been leveraging these resources productively in support of breeding (Schafleitner et al., 2022; van Etten et al., 2023). At the time of writing this publication, WorldVeg hosts a genebank containing 60,000 accessions from ∼400 vegetable species (World Vegetable Center, 2023). However, WorldVeg’s genetic resources for NUS are not heavily tapped by private seed companies, and formal NUS seed systems remain under-developed in many countries (Adebooye et al., 2005; Muendo et al., 2004). Better integration of WorldVeg’s genebank with public breeding programs and existing seed systems could help ensure sustainable access to a diversity of improved NUS varieties from which producers and consumers can benefit (N’Danikou et al., 2022). Over the last decade, WorldVeg’s African traditional vegetables breeding program has leveraged gender-disaggregated participatory breeding approaches to identify product profiles and select promising breeding lines of amaranth, African eggplant and other traditional vegetables (Dinssa et al., 2016, 2022), but this has been conducted in a limited number of locations. Surprisingly no or very few gender-segregated preference studies can be found for these traditional vegetables (summarized in Christinck et al., 2017; Weltzien et al., 2019). Evidence of gender-based differences in consumer preferences, demand, and willingness-to-pay for NUS is also scanty (Gido et al., 2017; Odendo et al., 2020; Senyolo et al., 2014; Wanyama et al., 2023).

This knowledge gap could be a crucial oversight given the gender dynamics of NUS production and marketing. Women are often heavily engaged in production of NUS in rural areas, where they typically manage home gardens and prepare food for the household (Dinssa et al., 2016; Ojiewo et al., 2015). Commercial vegetable production, in contrast with subsistence production in home gardens, skews toward men (Wanyama et al., 2023; Weinberger & Pichop, 2009).

However, women are primary marketers of NUS, even in cases where men are the primary producers (Dinssa et al., 2016; Fischer et al., 2020; Weinberger & Pichop, 2009). As such, understanding men’s and women’s preferences as producers, marketers, and consumers of NUS is critical and should be incorporated into analysis of market segments.

In this context, expanded participatory breeding research is critical to ensure that promising accessions selected from gene banks hosting NUS, and any improved varieties developed through them, respond to the real-world needs, constraints, and priorities of farmers and consumers (Schafleitner et al., 2022; van Etten et al., 2023; Van Zonneveld et al., 2023).

Participatory research also offers opportunities to explore diversity considerations and market segmentation, i.e., how gender, socioeconomic status, and intended product end-uses might contribute to variation in trait and varietal preferences. We used the tricot approach for on-farm testing, which allows evaluation of a collection of varieties for multiple traits across many women and men farmers and locations. It is this property that makes it possible to detect differential preferences across segments and understand how preferences differ among gender. In this process, we also sought to build a model for demand-driven participatory breeding that can be applied to other NUS as well as staple crops, supporting the expansion of local seed enterprises’ engagement in NUS seed systems as well as farmers’ access to quality NUS seed.

We used leafy amaranth as an example case to examine farmer preferences for NUS in different countries and identify market segments. Amaranth (*Amaranthus* spp.) is among the most commonly recognized traditional African vegetables, typically grown at small scale and often in home gardens (Ochieng et al., 2019). Economically important species include *A. cruentus, A. hypochondriacus, A. hybridus, A. dubius* and *A. caudatus* (Dinssa et al., 2016). Although amaranth originates as a grain crop in the Americas, it is consumed primarily as a leafy vegetable in Africa, with demand for grain building (van Zonneveld et al., 2021). Leaf nutrient content may vary with species and genotype. Most species constitute a good source of protein and calcium (particularly the grain), Vitamin C, zinc, magnesium, and other minerals (Kachiguma et al., 2015; Kamga et al., 2013). Although WorldVeg seed kit distributions have helped disseminate improved varieties (Stoilova et al., 2019; Wanyama et al., 2023), access to quality amaranth seed remains a challenge, as there is not yet a wide diversity of improved amaranth seed varieties in many markets (Cernansky, 2015; Kansiime et al., 2018; Onim & Mwaniki, 2008).

## Materials and Methods

### Trial design and variety evaluation

On-farm citizen science trials were conducted to enable participatory amaranth variety testing across a range of agroecological (humid coastal and drylands), socioeconomic (urban and peri-urban settings), and societal (cultural settings and gender) contexts. Trials were based on the triadic comparison of technologies (tricot) approach, in which a large number of farmer-managed plots are established on which individual farmers host random sets of three out of the full set of varieties and evaluate each of the three varieties at multiple stages in the growing season (van Etten et al., 2019, 2020). This design is particularly suited to the evaluation of varietal performance by many different farmers, which is essential for detecting differences in preference among types of producers. Unlike conventional participatory variety selection conducted mostly through researcher-managed trials, farmers participate individually. They assess varieties grown on their own fields with their own tools and inputs, using the labor to which they have access, in the context of their unique needs and constraints. In this regard, the tricot approach is relatively sensitive to gender and social inclusion (Voss et al., 2023).

Decentralized on-farm trials also mitigate some of the concerns that researcher-managed on-station and on-farm trials, including participatory variety selection through researcher-managed trials, are not representative of farmers’ actual growing conditions and are poor predictors of farmer preferences (De Roo et al., 2017; de Sousa et al., 2021; Laajaj et al., 2020; Misiko, 2013).

Amaranth trials for this study were conducted with 794 farmers (Figure 1) in the Atlantic, Oueme, and Alibori regions of Benin, and with 969 farmers in the Bougouni, Sikasso, and Koulikoro regions in Mali, from 2021-2022. Trials were also conducted with 300 farmers in Mtwara and Lindi regions in Tanzania in 2022. In all three countries, trials used a balanced incomplete block design under which host farmers received three of fourteen amaranth genotypes and promising accessions drawn from the WorldVeg genebank (Table 1). Each farmer received 2 g of seed per variety (6 g total), with variety names coded A, B, and C. Plot sizes (5 × 2 m in Mali and Tanzania, 6 × 1.2 m in Benin) and plant spacing (60 × 40 cm in Mali, 20 × 20 cm in Benin, 15 × 15 cm in Tanzania) were recommended but not strictly enforced. Farmers were permitted to practice their preferred management so long as it was consistent across their three plots.

**Figure 1.**
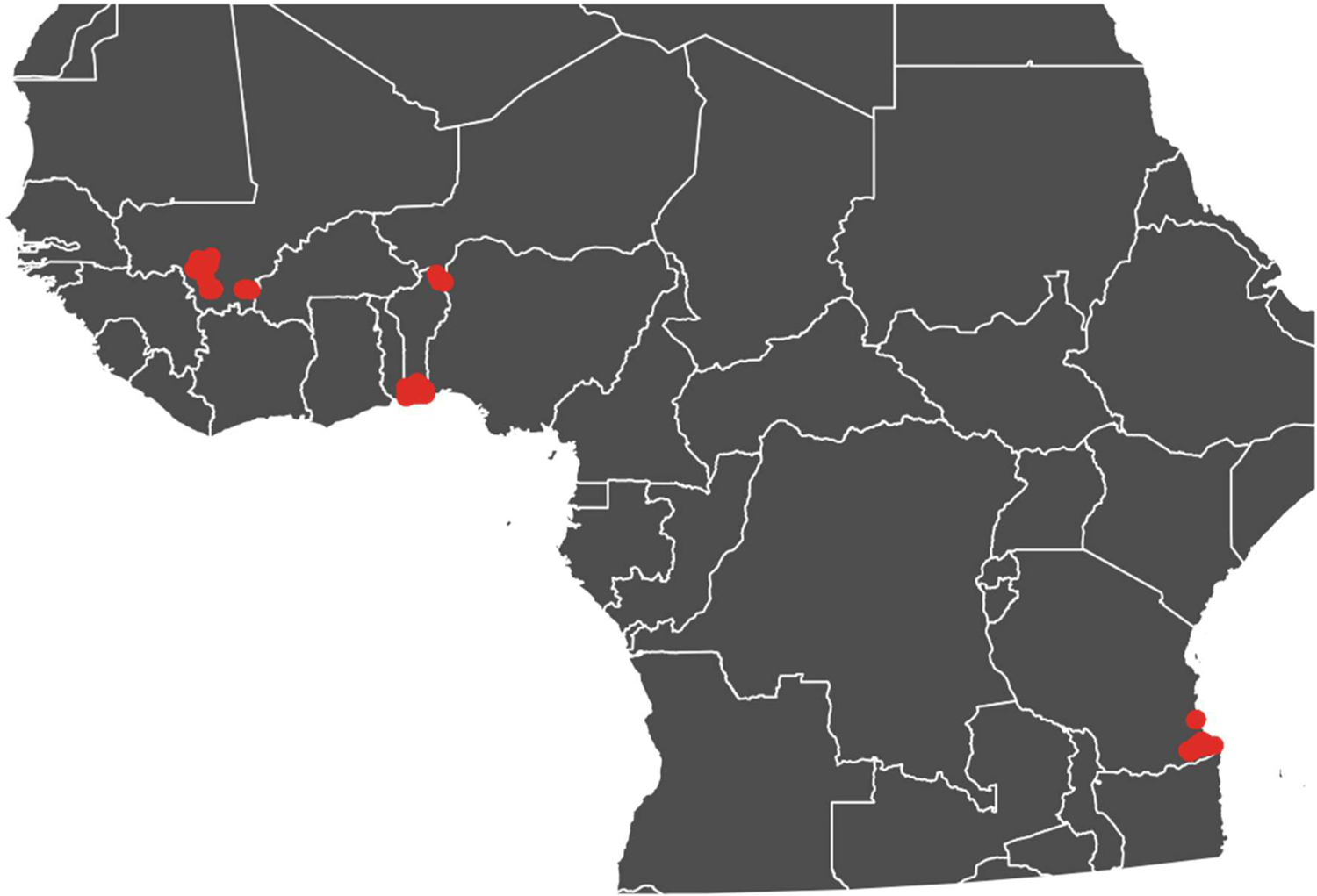
Centroids (red dots) of amaranth on-farm trials in Benin, Mali and Tanzania.

**Table 1.**
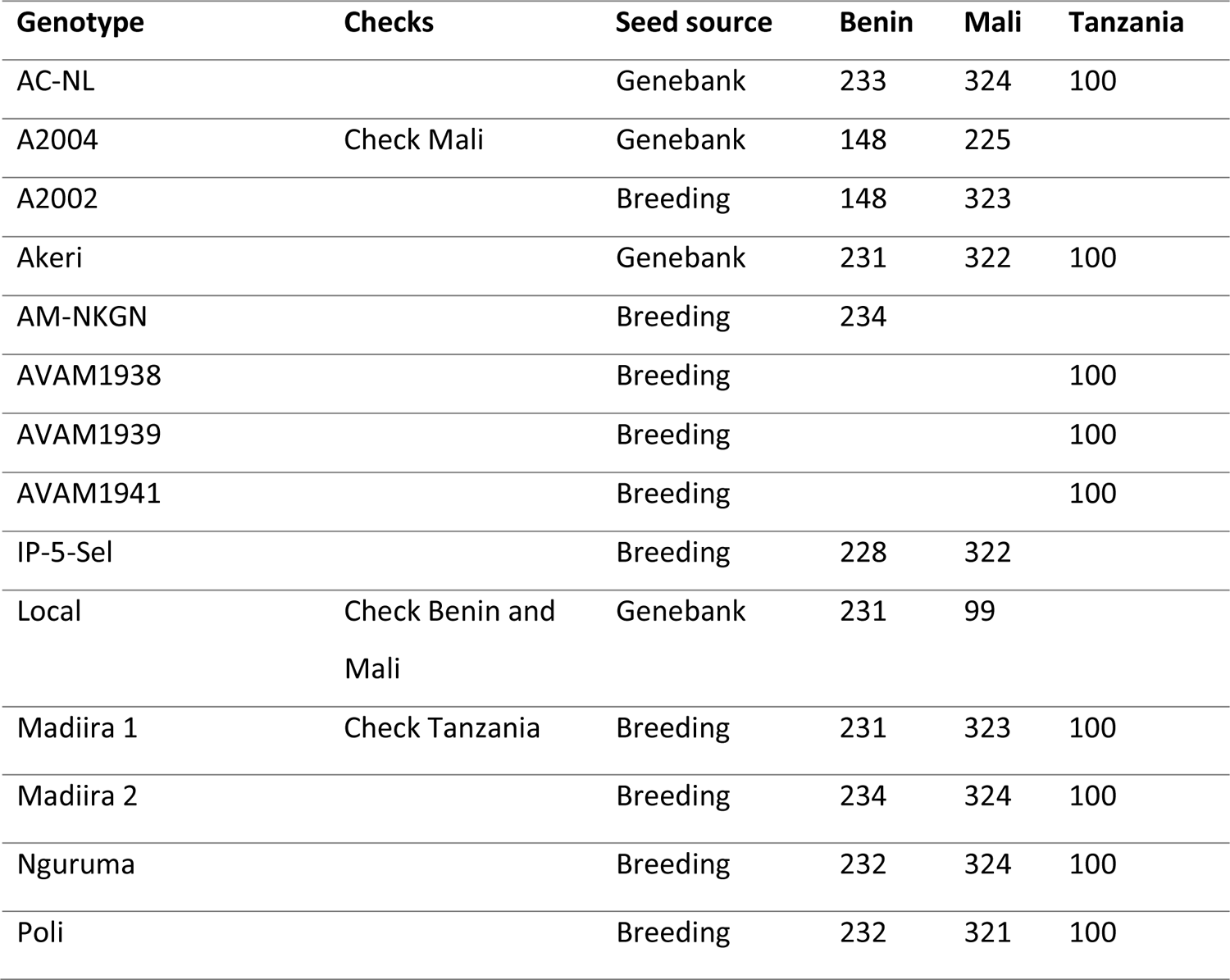
Varieties distributed in trials, and number of farmers per country.

| **Genotype** | **Checks** | **Seed source** | **Benin** | **Mali** | **Tanzania** |
| --- | --- | --- | --- | --- | --- |
| AC-NL |  | Genebank | 233 | 324 | 100 |
| A2004 | Check Mali | Genebank | 148 | 225 |  |
| A2002 |  | Breeding | 148 | 323 |  |
| Akeri |  | Genebank | 231 | 322 | 100 |
| AM-NKGN |  | Breeding | 234 |  |  |
| AVAM1938 |  | Breeding |  |  | 100 |
| AVAM1939 |  | Breeding |  |  | 100 |
| AVAM1941 |  | Breeding |  |  | 100 |
| IP-5-Sel |  | Breeding | 228 | 322 |  |
| Local | Check Benin and Mali | Genebank | 231 | 99 |  |
| Madiira 1 | Check Tanzania | Breeding | 231 | 323 | 100 |
| Madiira 2 |  | Breeding | 234 | 324 | 100 |
| Nguruma |  | Breeding | 232 | 324 | 100 |
| Poli |  | Breeding | 232 | 321 | 100 |

Farmers evaluated their three varieties regularly throughout the season, including four harvest periods. Evaluations involved ranking varieties’ overall performance and specific agronomic and end-use traits: germination, vigor, plant survival, pest tolerance, disease resistance, drought and flood tolerance, plant height, branching, yield, leaf size, marketability and taste (as consumers of their own products). Socio-economic data on the host farm and farmers, including information on product sales practices and seed acquisition, were also collected.

### Data analysis

We analyzed the tricot ranking data using the Plackett-Luce model (Luce, 1959; Plackett, 1975), recommended for analysis of on-farm tricot data (de Sousa et al., 2021). This model produces, scaleless, quantitative estimates of individual varietal performance for different traits, reflecting the probability of each variety of outperforming all other varieties in the tested set. The model is implemented in R using the package PlackettLuce (Turner et al., 2020) and extended with model-based recursive partitioning, which produces Plackett-Luce trees (Zeileis et al., 2008). We report probabilities of outperforming all other items in the set as log *worth* estimates. The data were processed using the R packages *ClimMobTools* (de Sousa & van Etten, 2024) and *gosset* (de Sousa et al., 2023). Due to the large number of traits assessed (10), some over multiple growth stages in the season (up to 6 data collection moments), we used Kendall Tau partial correlation to identify the traits most closely associated with farmers’ overall preference for the tested varieties. Traits were selected using a backward selection approach, starting with the full set of traits and iteratively removing the traits with least correlation to overall preference until no uncorrelated traits remain (p > 0.05). These traits were used to perform the likelihood-ratio test (described below), and the principal component analysis with the Plackett-Luce coefficients obtained for each trait.

To identify any potential farmers’ segments, we applied a cluster analysis using the farmers’ socioeconomic data (analyzed independently from the variety performance rankings). This was done following existing producer segmentation studies (Hammond et al. 2020; Kilwinger et al. 2021). We used covariates relevant to producer preferences identified in past studies and reflective of grower and end-user requirements in seed product market segments (Donovan et al., 2022). The variables used were gender, age, years of experience growing the amaranth, distance to markets, gender of who controls the production, gender of who controls the selling, and household income share from amaranth production (Table 2, Table 3). Segmentation was performed in R using the package cluster (Maechler et al., 2023). The categorical variables were converted to factors, and dissimilarities were computed using the daisy function. Numeric variables were standardized, and Euclidean distances were calculated. The resulting distance matrices underwent hierarchical clustering, and optimal clusters were determined via the cutree function. Subsequent refinement and validation of clusters were conducted, leading to the identification of four distinct clusters (segments). After the definition of segments in R, we used a Large Language Model approach to describe the main characteristics of each segment using the full set of covariates. The descriptions were checked and refined afterwards to prevent hallucinations, when incorrect or misleading results are generated.

**Table 2.**
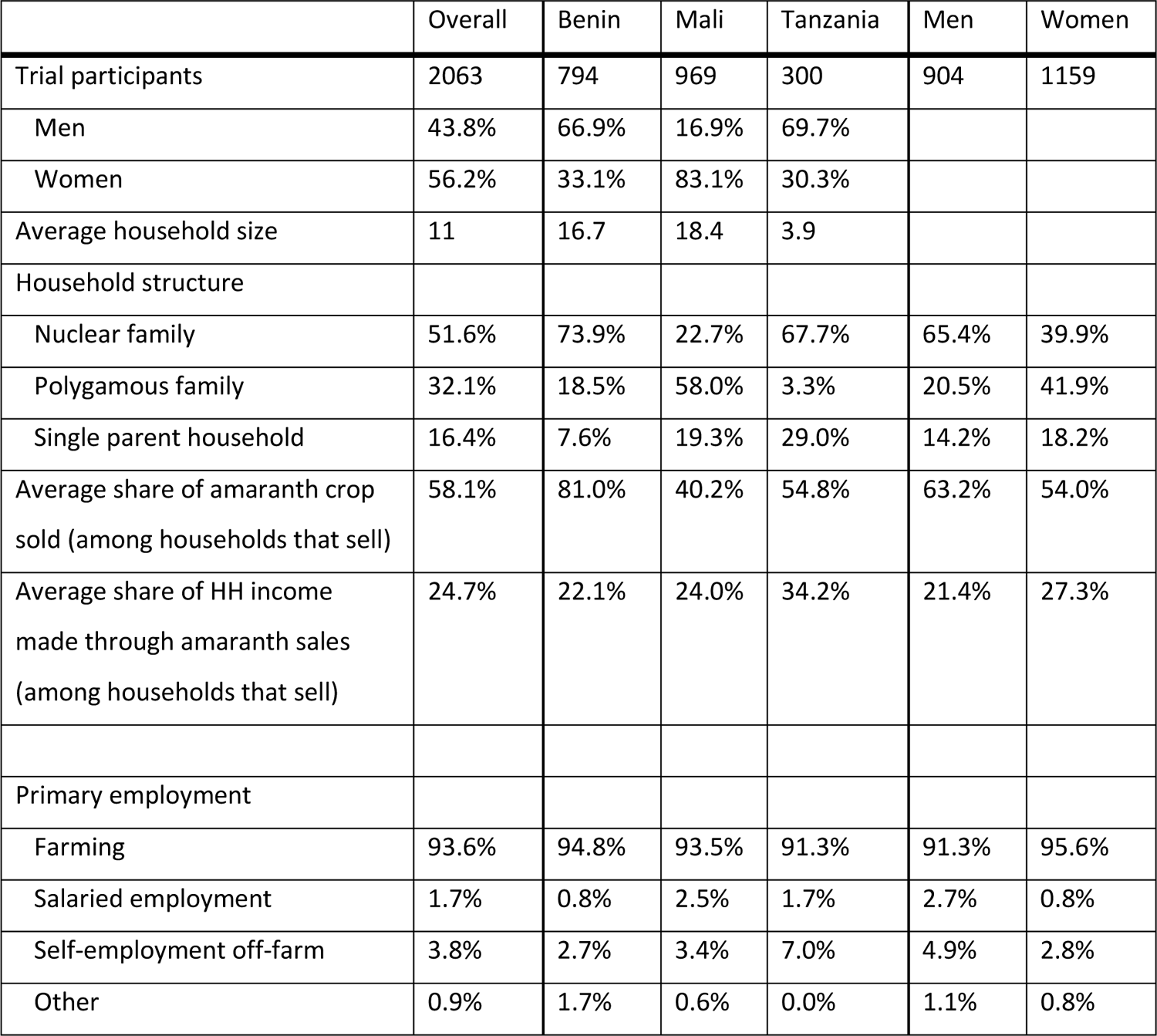
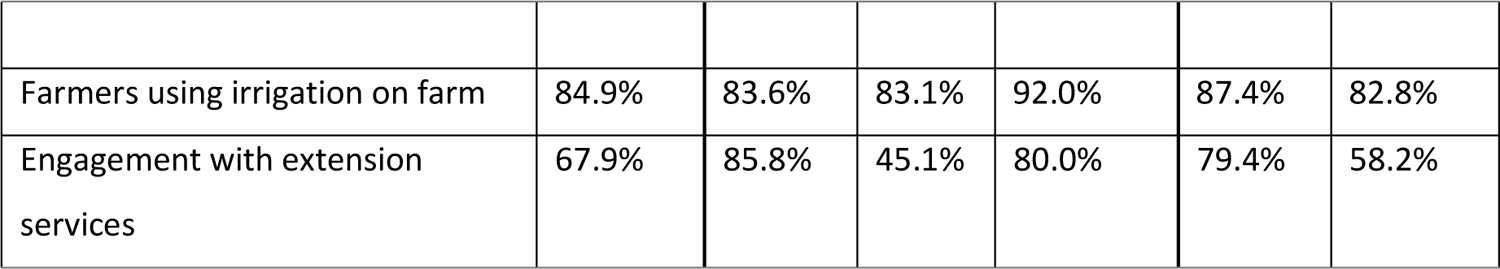
Characteristics of trial participants and their households/farms.

**Table 3.** Household-level control over amaranth production, sales, and seed exchange, as reported by the trial participants.

|  | Overall | Benin | Mali | Tanzania |
| --- | --- | --- | --- | --- |
| Control over amaranth production |  |  |  |  |
| Man | 40.0% | 58.9% | 13.2% | 60.0% |
| Woman | 48.2% | 29.3% | 75.1% | 27.7% |
| Both | 8.4% | 4.0% | 11.4% | 11.0% |
| Other | 3.4% | 7.8% | 0.3% | 1.3% |
| Control over amaranth sales |  |  |  |  |
| Man | 34.4% | 50.8% | 7.5% | 59.0% |
| Woman | 52.0% | 32.4% | 81.4% | 28.7% |
| Both | 7.8% | 4.7% | 9.5% | 10.7% |
| Other | 5.8% | 12.1% | 1.7% | 1.7% |
| Control over amaranth seed exchange |  |  |  |  |
| Man | 39.8% | 56.2% | 14.4% | 61.3% |
| Woman | 46.3% | 28.5% | 71.1% | 28.0% |
| Both | 10.1% | 5.5% | 14.4% | 9.3% |
| Other | 3.9% | 9.8% | 0.2% | 1.3% |

We then used a likelihood-ratio test to assess whether varietal rankings for key traits retained by the Kendall partial correlation differed significantly between different segments. We used the function likelihood_ratio available in the R package gosset. Briefly, the Plackett-Luce model is fitted using maximum likelihood, which allows the log likelihood for a single model fitted to full dataset to be compared to sum of log likelihoods for separate models fitted to pre-defined splits of the data, accounting for the increase in degrees of freedom in such a segmented model.

Finally, we performed a regret analysis using coefficients from the Plackett-Luce rankings of marketability. We used the function regret from the package gosset. Regret is a risk assessment analysis to support farmers’ diversification analysis. We present minimum regret, which is calculated by taking the summed squares of the distances of each variety to the best variety in each market segment and taking the square root of the resulting sum. It can be interpreted as the total distance to the ‘best variety’ in each group using the log-worth estimates. This measure is therefore more sensitive to differentiated preferences than the overall worth, which could be biased if one group has a very strong preference for a particular variety, which is less preferred by other segments.

## Results

### Sample characteristics

Registered trial participants’ demographic and socioeconomic characteristics are indicated in Table 2. Across countries, 56.2% of the trial participants were women, although men were disproportionately represented in Tanzania (69.7%) and Benin (66.9%) and under-represented in Mali (16.9%). This may reflect the higher degree of commercialization of amaranth in Benin and Tanzania. Roughly half of participating households were nuclear families, although the majority of women trial participants came from either polygamous or single parent households.

Farming was the dominant occupation across countries. Among the 84% of households that reported selling amaranth, on average 58% of amaranth produced was sold (81% in Benin). These sales contributed 25% of household income on average among households selling amaranth.

Table 3 shows the intrahousehold dynamics of amaranth production, sales, and seed exchange and suggests that women are slightly more engaged in amaranth activities overall, and especially sales. However, there is substantial variation in men’s and women’s engagement between countries. While amaranth production, marketing, and seed exchange were reportedly primarily undertaken by men in Benin and Tanzania, women in Mali were said to hold disproportionate responsibility for amaranth. These differences are, at least in part, likely a result of the gender balance of the trials in the three countries, as both women and men were more likely to report themselves as having control over amaranth than to report that their partner has control.

### Overall trait and variety preferences

To understand farmer preferences, we first used Kendall-Tau correlations to identify key traits correlated with farmers’ overall preferences. This process showed that plant survival (both during the reproductive phase and at the third harvest), yield (primarily at later harvests), taste and leaf size (at final harvest), and marketability (at all harvest periods) were the traits that drove farmers’ overall preferences (Figure 2). This allowed us to focus on these traits as those most relevant to farmers’ overall choices.

**Figure 2.**
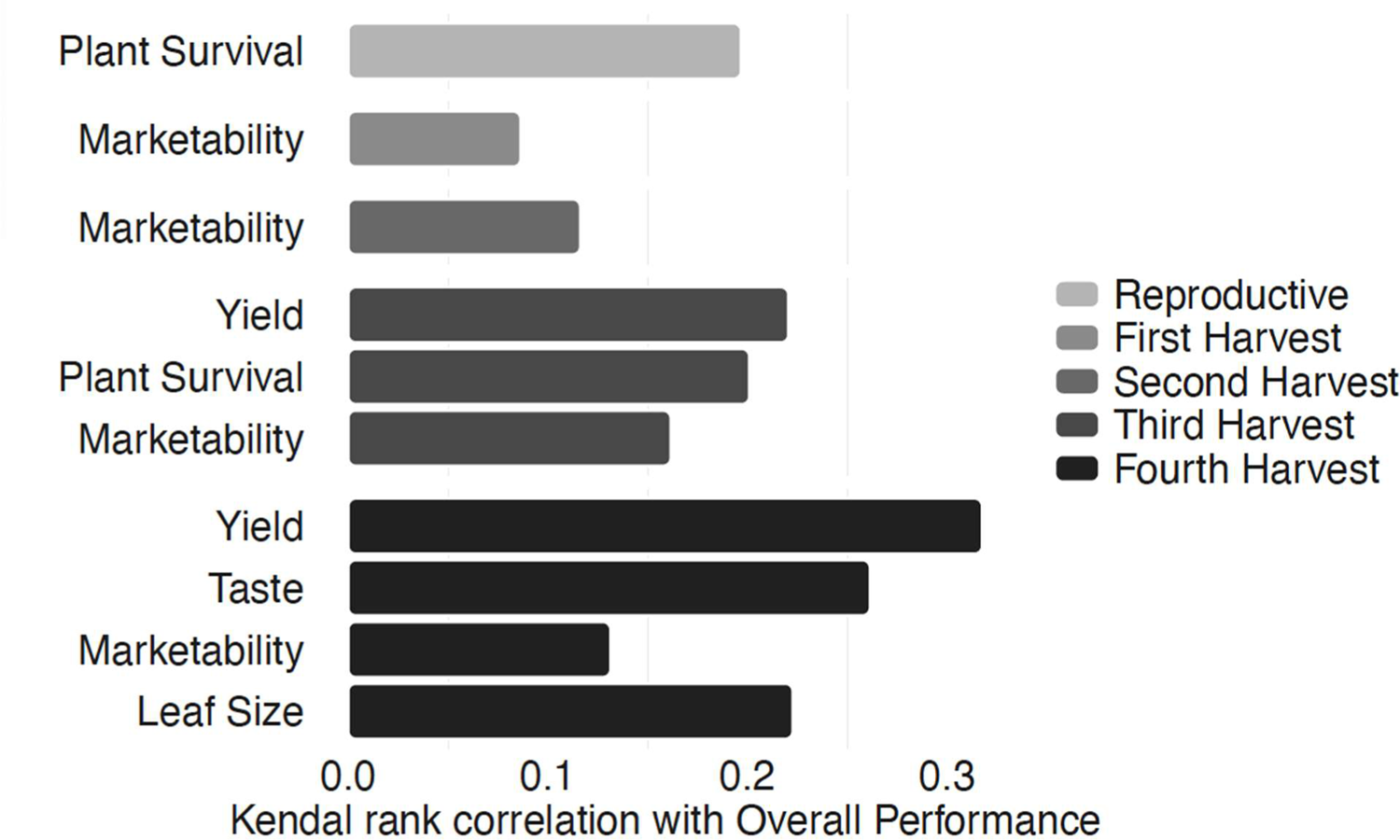
Among the many traits which farmers evaluated throughout the season, marketability, yield (especially at later harvests), taste, plant survival, and leaf size were most strongly correlated with farmers’ overall variety preferences.

In aggregate (Figure 3), farmer overall variety preferences skewed toward Akeri, Poli, and AM-NKGN, linked to their marketability, leaf size, yield, and taste at fourth harvest. These varieties were often reported to be preferred for use in subsequent seasons. A2004 emerged as another popular variety, driven by its marketability in the first three harvests, yield, and plant survival in the third harvest, but less by overall preference. However, this breakdown ignores any possible segmentation of farmer preferences.

**Figure 3.**
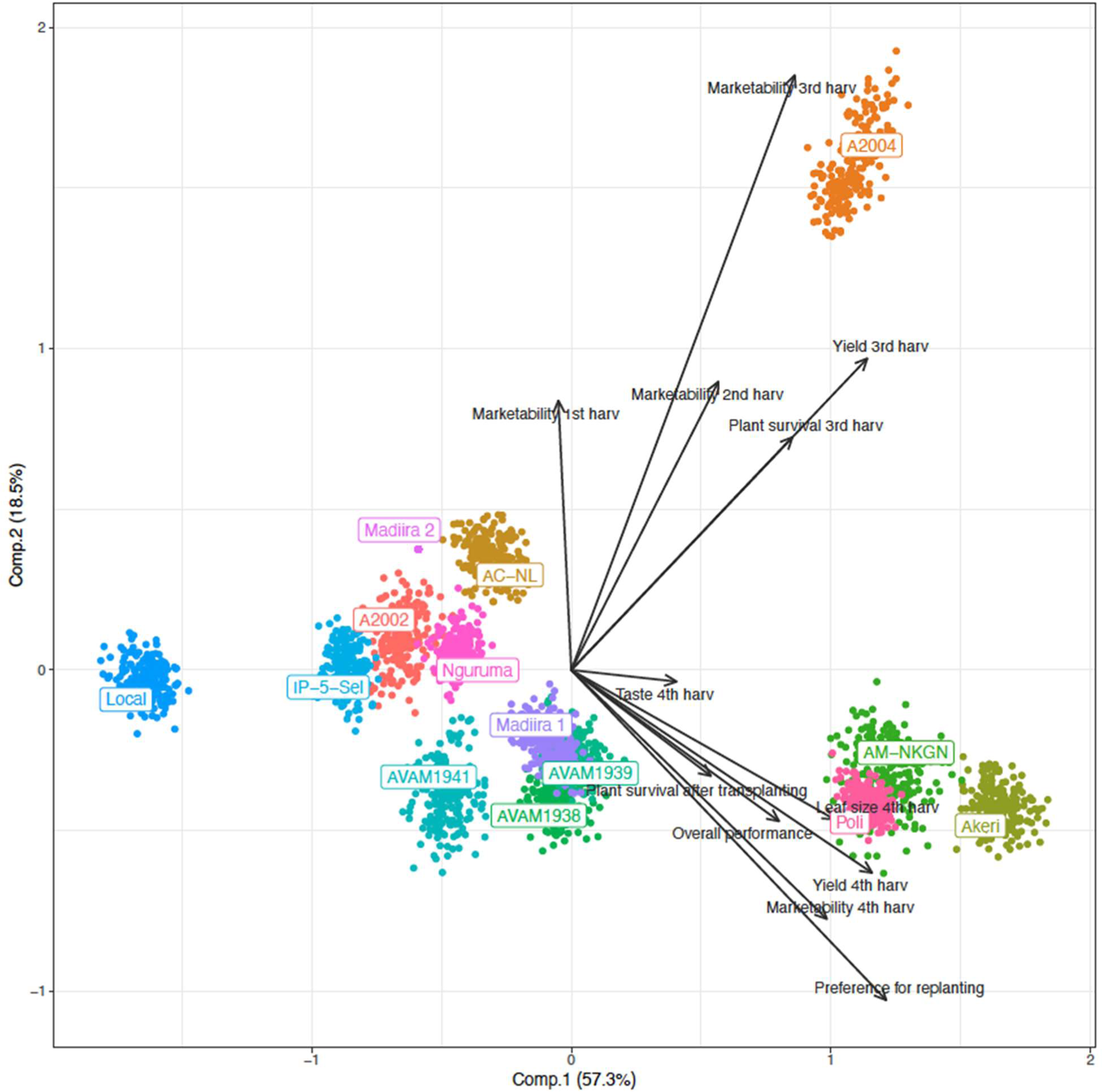
Principal components of Plackett-Luce model estimates (log-worth) on amaranth trait performance. Dots represent the performance (log-worth) of each amaranth variety ranked by farmers. Arrows represent the paths (correlation) of varieties and the main traits retained after backward selection.

### Variation in farmer preferences

To consider variation in farmer preferences, we undertook a segmentation process that produced four farmer segments, listed in Table 4. These segments are largely distinguished by gendered control over amaranth, income generated, and experience in amaranth farming. The “Older Women Generalists” segment represents women who have significant control over both the sale and production of amaranth. They earn only a moderate share of income from amaranth and have relatively less experience in amaranth farming. “Young Women Specialists” includes younger women who were highly involved in both the production and sale of amaranth. They boasted the highest income share from amaranth and substantial experience cultivating it, indicating a greater degree of specialization in amaranth. “Older Men Generalists” are predominantly men with considerable experience in amaranth farming. They have control over both the production and sale of amaranth but a lower income share from the crop. The “Young Men Specialists” segment represents younger men with the highest average experience cultivating amaranth and a significant income share from amaranth. They controlled both the production and sale.

**Table 4.**
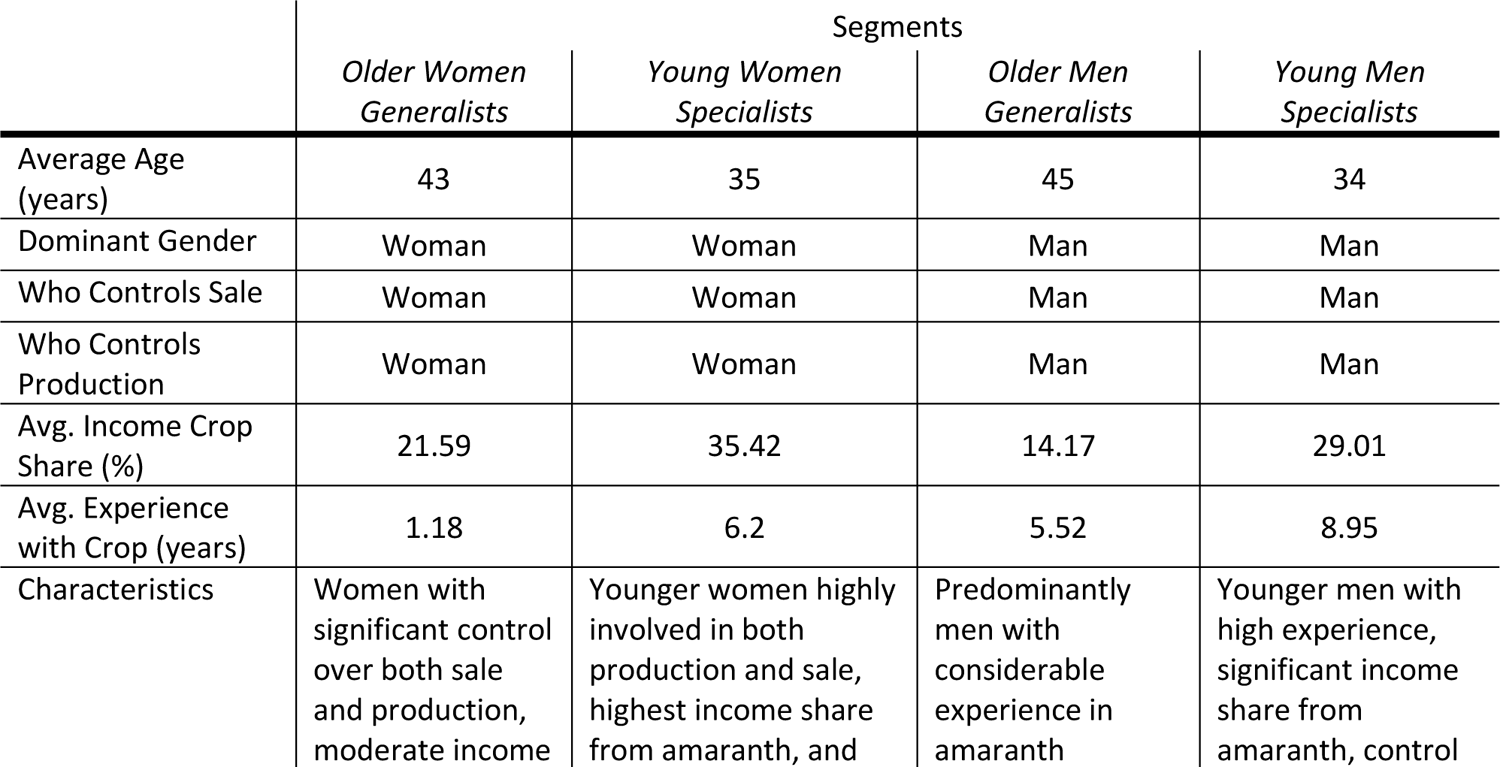
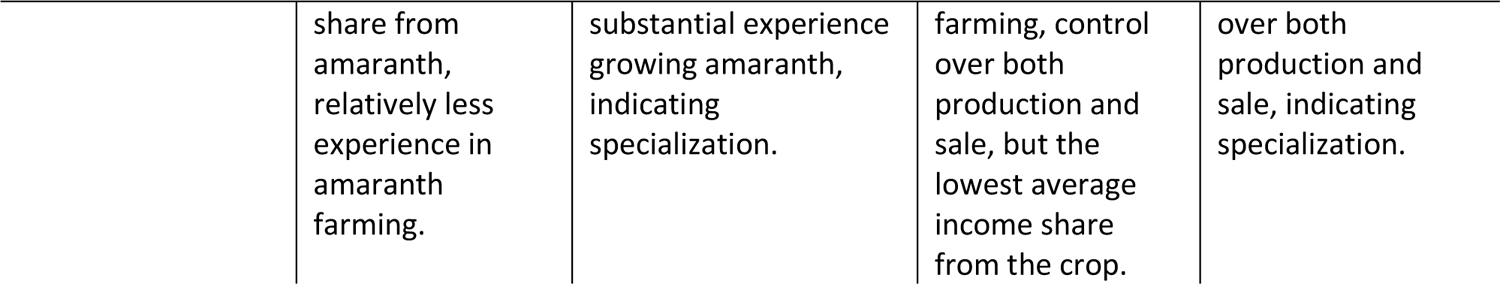
Demographic and socioeconomic characteristics of farmers segments in amaranth production in Benin, Mali, and Tanzania.

To validate these groups, we conducted a log-likelihood ratio test. Table 5 indicates whether each segment generated statistically different rankings on the key traits retained by the Kendall partial correlation. Other than traits ranked at the fourth harvest (when a smaller number of observations were recorded), all key traits’ rankings were distinguished across the four segments.

**Table 5.**
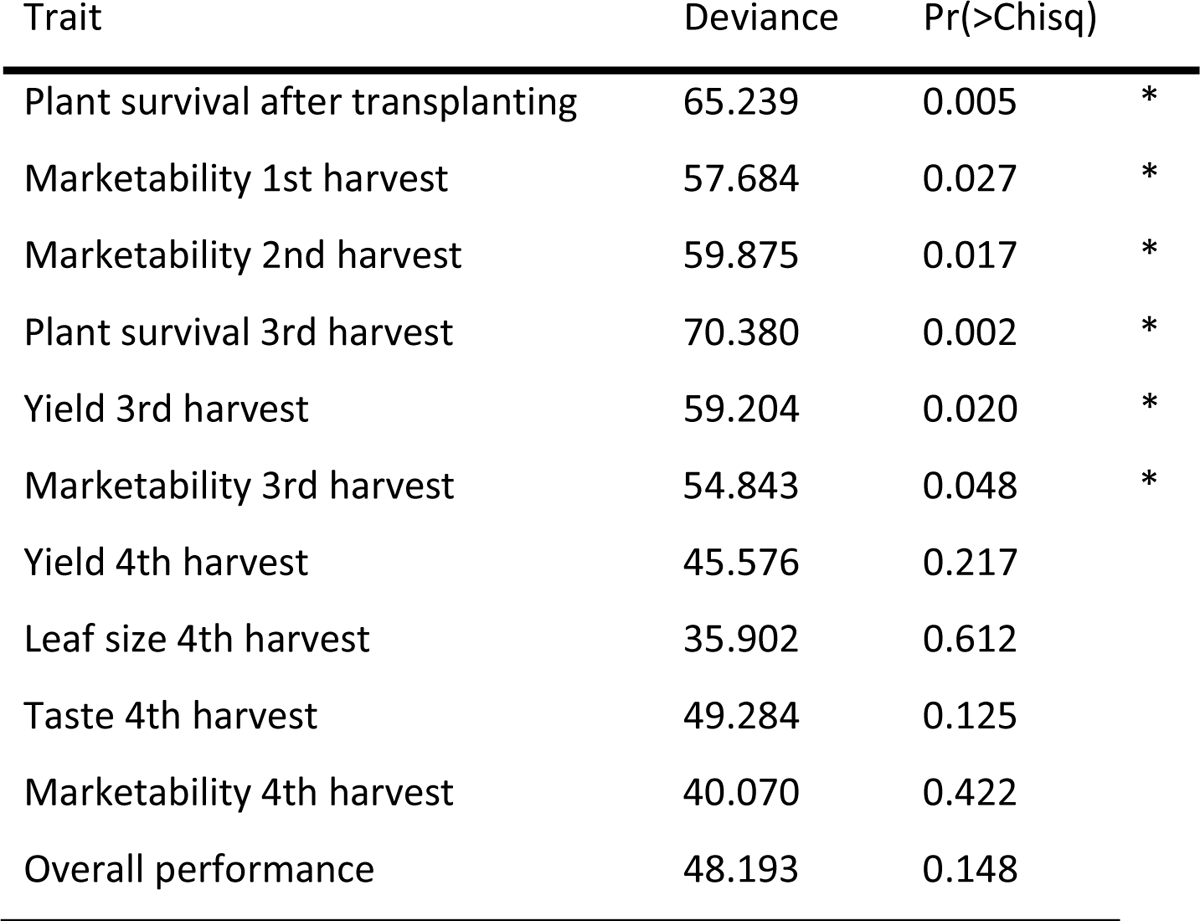
Log-likelihood ratio test estimates for the main traits assessed by farmers within segments.

| Trait | Deviance | Pr(>Chisq) |  |
| --- | --- | --- | --- |
| Plant survival after transplanting | 65.239 | 0.005 | * |
| Marketability 1st harvest | 57.684 | 0.027 | * |
| Marketability 2nd harvest | 59.875 | 0.017 | * |
| Plant survival 3rd harvest | 70.380 | 0.002 | * |
| Yield 3rd harvest | 59.204 | 0.020 | * |
| Marketability 3rd harvest | 54.843 | 0.048 | * |
| Yield 4th harvest | 45.576 | 0.217 |  |
| Leaf size 4th harvest | 35.902 | 0.612 |  |
| Taste 4th harvest | 49.284 | 0.125 |  |
| Marketability 4th harvest | 40.070 | 0.422 |  |
| Overall performance | 48.193 | 0.148 |  |

Figure 4, in contrast with Figure 3’s aggregated model, shows that trait and variety preferences differed substantially across the four farmer segments. For example, Older Women Generalists expressed a strong preference for Akeri, Poli, and AVAM1941, and consistent dislike of the other varieties (those in the western quadrants of Figure 4A). Young Women Specialists similarly identified five clearly preferred varieties (A2004, Akeri, AVAM1939, AM-NKGN, and Poli) and disliked the remainder. Older Men Generalists’ variety preference model shows a much more even spread of arrows and variety clusters, indicating little agreement about optimal varieties. Young Men Specialists’ model indicates strong preferences for A2002 and A2004 varieties driven by their marketability, while a large number of other varieties (Madiira 1, Nguruma, AC-NL, and Akeri) were preferred for other traits such as yield, leaf size, and plant survival.

**Figure 4.**
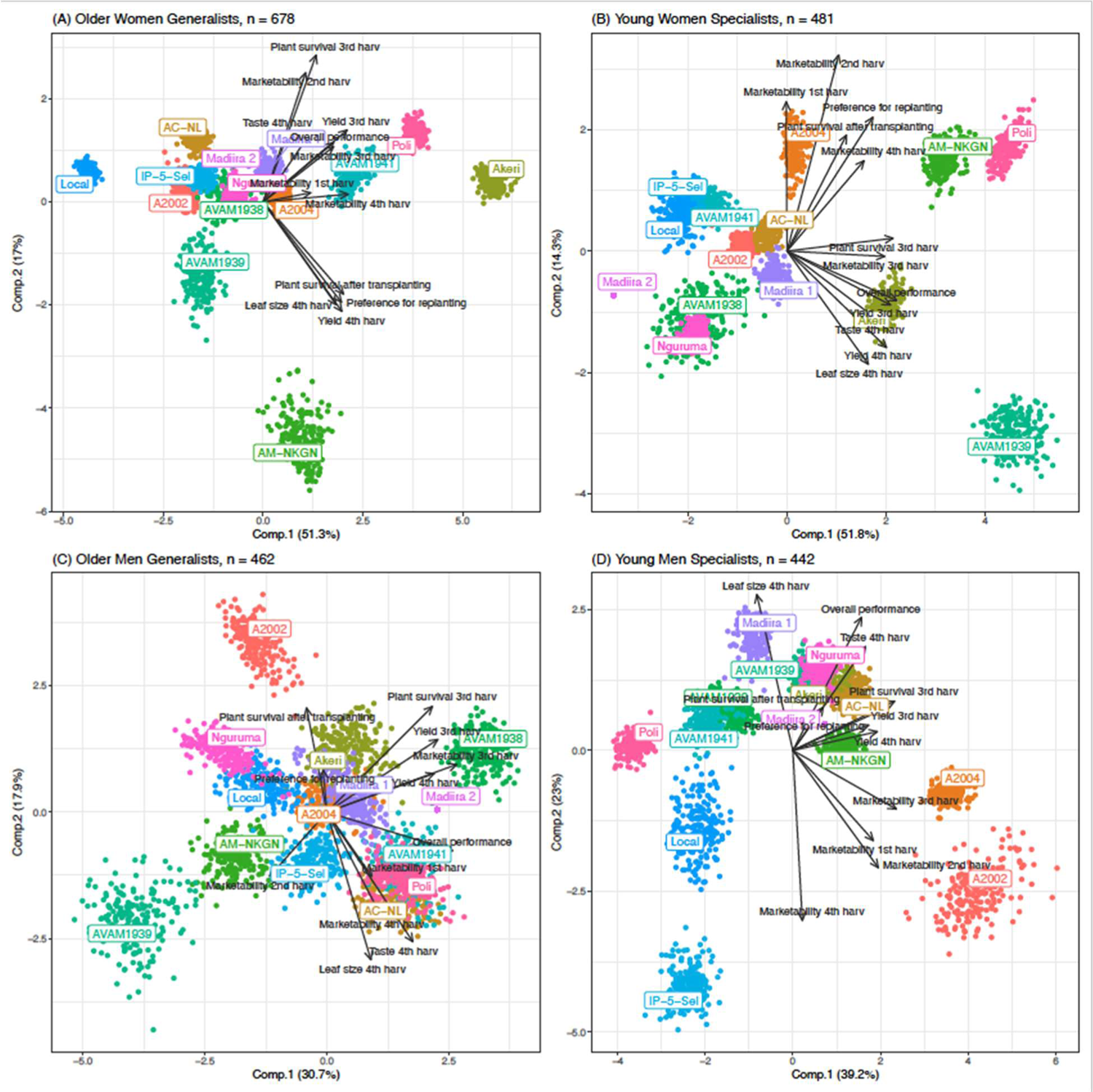
Principal components of Plackett-Luce model estimates (log-worth) on farmers’ segments. Dots represent the performance (log-worth) of each amaranth variety ranked by farmers. Arrows represent the paths (correlation) of varieties and the main traits retained after backward selection.

Finally, to provide a synthetic analysis to inform decision making, we present both worth and minimum regret values for each of the varieties (Table 6). This measure gives an indication of the ‘loss’ that would be perceived by the different segments compared to the variety that each group ranked as the best. Independent from the measure taken, worth or regret, A2004 would be acceptable to all groups. However, if a second variety is to be recommended, AC-NL could be chosen based on its overall high worth, but Poli would minimize regret. Choosing AC-NL would indeed mainly benefit Older Men Generalists, whereas Poli is preferred by the three other groups, and therefore a more balanced choice.

**Table 6.**
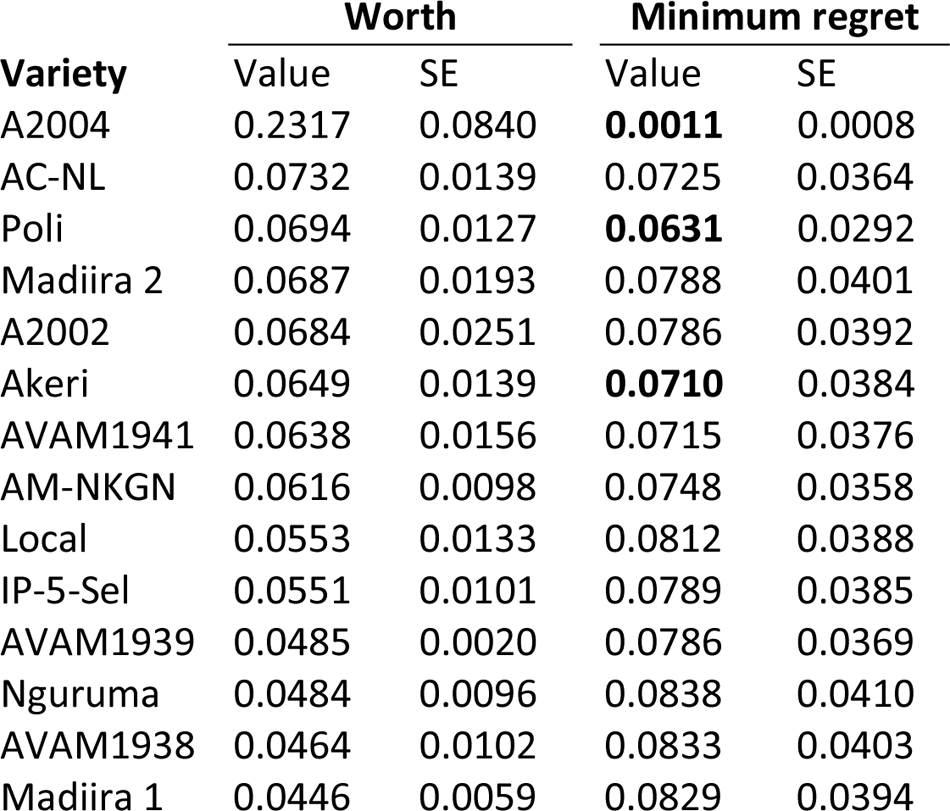
Average worth and minimum regret values and standard errors for the trait ‘marketability’ of evaluated amaranth varieties. Values in bold highlight the three varieties with smaller minimum regret.

## Discussion

### Implications for demand-driven amaranth breeding

These results provide useful information for public and private breeding programs of neglected vegetables and seed producers. First, they point to a distinct segmentation of amaranth farmers based on gendered control of amaranth production and sales, income generation, and experience producing amaranth. Variety preferences differed significantly across these segments, underscoring the heterogeneity of NUS producers and the value of segmentation on the basis of more than gender alone.

The results also provide specific insights into trait preferences to guide breeding programs. Farmers’ overall preferences for amaranth related primarily to yield, taste, plant survival from early to late stages of the season, leaf size, and marketability. These bear resemblance to priority traits documented in other studies in Tanzania (Adeniji & Aloyce, 2013; Dinssa et al., 2022). In a recent study in Tanzania, for instance, both women and men farmers ranked (1) fast growth habit (early biomass accumulation) plus quick recovery from repeat harvests, (2) marketability, and (3) ability to be harvested several times from the same planting material as the three most important traits (Dinssa et al., 2022). With these insights, and specific variety preferences in Figure 4, breeders can prioritize new crosses to meet current and future demand (Donovan et al., 2022).

Notably, although taste emerged as a key factor driving farmers’ preferences, it is not consistently included in breeding programs as a priority trait. Organoleptic properties’ historic exclusion from varietal testing and release processes are likely one reason for this (Afari-Sefa et al., 2012; Keatinge et al., 2015; Schreinemachers et al., 2021; Turner & Bishaw, 2016), and perhaps related to the limited attention paid to consumer preferences in many breeding programs (Thiele et al., 2020). Infamously, tomato breeding in the Netherlands had to drastically switch course and better respond to consumer preferences after the market collapsed due to tasteless tomatoes (Schouten et al., 2019). Increased attention to organoleptic properties may have particular relevance for gender-responsive and gender-intentional breeding and seed systems development, given evidence that women disproportionately prioritize these traits across crops (Weltzien et al., 2019). Breeding programs of neglected vegetables should account for consumer taste from the early stages of breeding programs, as WorldVeg’s amaranth breeding program has started doing (Dinssa et al., 2022). From a practical standpoint, feeding consumer preferences into breeding pipelines will require systematic assessment and participatory evaluation of organoleptic traits, their translation into measurable and breedable targets, and design of phenotypic assays.

### Gender implications

The results underscore, first, the relevance of gender considerations in amaranth breeding, with implications for wider breeding of neglected vegetables. The dynamics of intrahousehold control of amaranth production, sales, and seed exchange (Table 3) indicate that both women and men are (or at least perceive themselves to be) deeply involved in amaranth-related activities within households. This aligns with a previous study in Tanzania that found production activities for leafy vegetables to be shared, although they identified seed selection for vegetables to be largely men’s responsibility (Fischer et al., 2017). As such, understanding and appealing to both men and women’s needs, priorities, and constraints in breeding and seed system development are critical—especially when variety preferences differ, as found here. Further attention to the gender dynamics of seed selection and acquisition is also warranted to ensure men’s and women’s ability to equitably *access* and *benefit from* improved varieties.

Our results contrast with those from a more conventional, gender-disaggregated participatory varietal selection study of amaranth in Tanzania, where female and male farmers’ variety preferences were found to be similar (Dinssa et al., 2022). This discrepancy likely results in part from our use of a citizen science-based approach focused on understanding producer and consumer preferences grounded in men’s and women’s realities (Voss et al., 2023). Our analysis using market segments rather than gender-based disaggregation also illustrates how preferences may vary among women (and among men) according to their production orientation, experience, livelihood portfolio, and other factors. Such intersectional analysis of seed product market segments are likely to yield deeper insights into preferences than conventional gender-based disaggregation conducted by bringing farmers to centrally managed trials during a single moment in the crop cycle (Dinssa et al., 2022).

To illustrate this last point, it is especially interesting in this case that Older Men Generalists’ variety preferences are not well aligned with the other segments’ and are inconsistent. This may simply reflect disagreement within this segment, but more likely indicates that these farmers are not as certain as other farmer groups about which traits are desirable. This could result from a lack of expertise in the Older Men Generalists group, which is possible given that women are known to be disproportionately involved in amaranth production and marketing in many rural contexts, and that men could have overstated their own role in amaranth cultivation in this study. This possibility is concerning given that older men’s voices are often disproportionately elevated in decision-making, including around topics like breeding. Ensuring that the preferences of younger amaranth specialists and older women are adequately captured may be key to appropriately meeting current and future seed demand. We have demonstrated that using worth as a criterion that could lead to selecting a variety that is indeed only top-ranked by Older Men Generalists. We show that using minimum regret across segments as a decision-making criterion can lead to a more gender-sensitive selection that would benefit a larger and more diverse group of farmers.

### Supporting expanded production and consumption of NUS

The results of this study would, for perhaps the first time, enable seed companies and other seed producers to target specific market segments for NUS development, and specifically women and youth as these two groups are often trained and supported in NUS production. With more actors involved in the breeding and distribution of quality NUS seed, producers may be able to access more locally-adapted, climate-resilient, pest- and disease-tolerant, and nutritious varieties (Schreinemachers et al., 2021). While our findings can help expand breeding of neglected vegetables and improve seed access, efforts to improve the appeal of NUS varieties for producers must ultimately be paired with attention to consumer demand, value chain development, and policy changes (McMullin et al., 2021). Local knowledge around utilization of these crops, breeding in relation to consumer preferences, and improved post-harvest handling are all critical (Keatinge et al., 2015; Schreinemachers et al., 2018). There is also need for value chain development that offers greater potential for producers—and seed enterprises—to profit from NUS sales and NUS seed production (Onim & Mwaniki, 2008). This includes attention to postharvest processes and infrastructure to enable proper handling and storage of perishable vegetable products (Keatinge et al., 2015; Schreinemachers et al., 2021).

## Conclusion

In this study, we identified producer preferences for improved amaranth varieties and found variation across four distinct segments of farmers, which were differentiated by gender, income generation, and experience growing amaranth. We also identified the top traits of interest for farmers: plant survival, yield, leaf size, taste, and marketability drove farmers’ overall varietal preferences. Finally, we found evidence that perceptions of varieties’ marketability did not, for the most part, change over stages of the growth season.

The findings can help guide breeding programs and seed companies in expanding access to a suitable diversity of improved amaranth varieties, and specifically to reach women and youth. This study also provides a model for using available genebank accessions and participatory, demand-driven breeding approaches to inform development of improved NUS varieties, for which little breeding work has thus far been done. This is particularly timely because of the increased interest in breeding of neglected vegetables (Fredenberg et al., 2024). Our study can inform these and other initiatives on how citizen science can support demand-driven breeding of improved NUS varieties with higher yields, more climate resilience, and improved nutrition that respond to diverse market segments’ needs, priorities and constraints (van Etten et al., 2023; Van Zonneveld et al., 2023). Through this, public and private breeding institutions can support expanded production and consumption of NUS across Africa.

## Acknowledgements

This work was supported by the German Federal Ministry for Economic Cooperation and Development (BMZ) commissioned by the Deutsche Gesellschaft für Internationale Zusammenarbeit (GIZ) through the Fund International Agricultural Research (FIA), grant number: 81260859 to the project “Choose, Grow, Thrive: Using citizen science in expanding West Africa’s food basket with African vegetables to tackle malnutrition (BMZ-CGT)”; the Swiss Agency for Development and Cooperation (SDC) through the Consumption of Resilient Orphan Crops & Products for Healthier Diets (CROPS4HD) project; and the Taiwan Africa Vegetable Initiative. Funding for Africa’s vegetable genebank is provided by other long-term strategic donors to the World Vegetable Center: Republic of China (Taiwan), United States Agency for International Development (USAID), Australian Centre for International Agricultural Research (ACIAR), United Kingdom, Thailand, Philippines, Korea, and Japan. Additional research time for KdS and MAL was provided through the 1000FARMS project (INV-031561) funded by the Bill & Melinda Gates Foundation.

## Author Contribution

SN, KdS, and MvZ conceptualized the research; MvZ, SN and AR acquired funds; SN, FD, AS, LA, ECL, ACH and AD coordinated data collection; KdS, MAL and JvE performed analysis and developed figures; RCV and KdS developed the first draft. All authors reviewed and contributed to first and later drafts.

## Data Availability Statement

Data and code are available on GitHub https://github.com/AgrDataSci/amaranth-worldveg/

